# Dissociation of reliability, heritability, and predictivity in coarse- and fine-scale functional connectomes during development

**DOI:** 10.1101/2022.05.24.493295

**Authors:** Erica L. Busch, Kristina M. Rapuano, Kevin M. Anderson, Monica D. Rosenberg, Richard Watts, BJ Casey, James V. Haxby, Ma Feilong

## Abstract

The functional connectome supports information transmission through the brain at various spatial scales, from exchange between broad cortical regions to finer–scale, vertex–wise connections that underlie specific information processing mechanisms. In adults, while both the coarse- and fine-scale functional connectomes predict cognition, the fine-scale can predict up to twice the variance as the coarse-scale functional connectome. Yet, past brain-wide association studies, particularly using large developmental samples, focus on the coarse connectome to understand the neural underpinnings of individual differences in cognition. Using a large cohort of children (age 9 – 10 years; *n* = 1,115 individuals, both sexes, 50% female, including 170 monozygotic and 219 dizygotic twin pairs and 337 unrelated individuals), we examine the reliability, heritability, and behavioral relevance of resting-state functional connectivity computed at different spatial scales. We use connectivity hyperalignment to improve access to reliable fine-scale (vertex–wise) connectivity information and compare the fine-scale connectome with the traditional parcel–wise (coarse scale) functional connectomes. Though individual differences in the fine-scale connectome are more reliable than those in the coarse-scale, they are less heritable. Further, the alignment and scale of connectomes influence their ability to predict behavior, whereby some cognitive traits are equally well predicted by both connectome scales, but other, less heritable cognitive traits are better predicted by the fine-scale connectome. Together, our findings suggest there are dissociable individual differences in information processing represented at different scales of the functional connectome which, in turn, have distinct implications for heritability and cognition.

**Significance statement:** Years of human magnetic resonance imaging (MRI) research demonstrate that individual variability in resting-state functional connectivity relates to genetics and cognition. However, the various spatial scales where individual differences in connectivity could occur have yet to be considered in childhood brain– behavior association studies. Here, we use novel machine learning approaches to examine the reliability, heritability, and behavioral relevance of different spatial scales of the resting-state functional connectome during childhood. We show that broad features of the connectome are strongly related to heritability, whereas fine details are more reliable and strongly associated with neurocognitive performance. These data indicate that reliable, heritable, and behaviorally–relevant individual differences exist at dissociable scales of the functional connectome.

## Introduction

The development of the functional connectome is an interplay of genetics (Glahn et al., 2010; Jansen et al., 2015; Miranda-Dominguez et al., 2018) and experience (Blokland et al., 2012; Yang et al., 2016). Functional connections at various spatial scales account for related yet distinct information processing (Haak et al., 2018; Haxby et al., 2020). While variation in functional connectivity (FC) predicts individual differences in memory (Lin et al., 2021), attention (Rosenberg et al., 2016), and fluid intelligence (Finn et al., 2015), among other traits (Dosenbach et al., 2010; Cui et al., 2020; Sripada et al., 2020; Feilong et al., 2021; Chen et al., 2022), how genetics and experience guide the development of different scales of FC remains unclear. Here, we estimate the heritability, reliability, and predictivity of resting-state FC (RSFC) at different levels of granularity in children. Our data suggest that the spatial scale of the functional connectome dissociates effects of experience-based RSFC features from heritable influences.

Past studies revealed genetic control over brain morphology (Thompson et al., 2001; Peper et al., 2009; Blokland et al., 2012; Brouwer et al., 2014; Elliott et al., 2018) and RSFC (Glahn et al., 2010; Fu et al., 2015; Colclough et al., 2017; Ge et al., 2017; Miranda-Dominguez et al., 2018; Anderson et al., 2021; Barber et al., 2021). Genetic influences on brain function are associated with cognition, increase with age, and interact with the environment to sculpt idiosyncratic functional connectomes (Lenroot et al., 2009; Brouwer et al., 2014; Schmitt et al., 2014; Jansen et al., 2015). The development of brain-behavior associations is often interpreted as a passive maturation, where genetics determine brain development which determines behavior. However, more recent frameworks suggest an interactive specialization of the developing brain that involves bi-directional interactions between genes, brain regions and networks, and psychological function across age and experience (Johnson, 2011). The interactions guiding and shaping functional brain-behavior associations in development likely vary at different spatial scales of functional connectivity.

Whole-brain RSFC is commonly calculated by parcellating the cortex and averaging over vertices within a parcel, then correlating the timeseries between pairs of parcels. Parcellations mitigate high-dimensionality, measurement instabilities, and idiosyncratic functional topographies by coarsely aligning fMRI signals, and have been used in prior developmental brain–behavior association studies (Sripada et al., 2020; Chen et al., 2022; Marek et al., 2022). Parcellations assume the same regional boundaries exist across brains, regardless of individual or developmental differences (Bandettini et al., 2022). Individual– defined parcellations better capture individualized functional topography (Glasser et al., 2016; Kong et al., 2019, 2021, 2023), yet cover large surface areas and collapse over local, fine-scale information processing. A useful alternative, functional alignment maximizes inter-subject correspondence while resolving fine-scale anatomical variation. Meaningful individual differences in fine-scale signals are most apparent after resolving functional topographies with methods like hyperalignment (Haxby et al., 2011, 2020). Instead of modeling functional topographies in anatomical dimensions, hyperalignment models functional information as high–dimensional pattern vectors and optimally aligns them to a common information space. The dimensions of this space represent shared functional patterns, retaining granularity while resolving fine-scale mismatches. Using hyperalignment to model the developing fine-scale functional connectome yields two main advantages. First, hyperalignment improves between–subject correspondence (Guntupalli et al., 2016, 2018; Busch et al., 2021) and highlights reliable, behaviorally– relevant idiosyncrasies (Feilong et al., 2018, 2021). Second, hyperalignment can disentangle anatomical from functional topographies to reveal how fine-scale RSFC relates to genetics and cognition, independent of anatomical idiosyncrasies.

To date, genetic contributions to fine-scale FC remain to be quantified, as well as their reliability and relation to developing cognition. Here, we ask: Does accessing the fine-scale connectome tighten the link between FC, cognition, and genetics shown in coarse-scale connectivity? Or is there a dissociation of genetic control over these two scales? We use connectivity hyperalignment (Guntupalli et al., 2018) to access and dissociate coarse- and fine-scale RSFC in 1,115 children from the ABCD Study®. We leverage machine learning approaches to understand how connectome granularity relates to reliability, heritability, and cognition in children.

## Materials and methods

### The ABCD Study

We considered a subset of data from the 11,875 children included in the ABCD Study data release 2.0.1 (Casey et al., 2018). The ABCD Study is a longitudinal study with 21 sites around the United States and aims to characterize cognitive and neural development with measures of neurocognition, physical and mental health, social and emotional function, and culture and environment. The ABCD Study obtained centralized institutional review board (IRB) approval from the University of California, San Diego, and each site obtained local IRB approval. Ethical regulations were followed during data collection and analysis. Parents or caregivers provided written informed consent, and children gave written assent. Four leading twin research centers at the University of Minnesota, Virginia Commonwealth University, University of Colorado-Boulder, and Washington University in St. Louis comprise the ABCD Twin Hub. Each site enrolled approximately 200 same-sex monozygotic (MZ) or dizygotic (DZ) twin pairs as well as singletons (Iacono et al., 2017). Their inclusion in the ABCD study affords unique access to the causal interrelation between genetics, environment, brain function, and cognition during development (Blokland et al., 2012; Jansen et al., 2015; Iacono et al., 2017)

The present study used imaging and behavioral data from a subset of *n* = 1,115 subjects from the original 11,875 subjects. Included subjects were enrolled in the University of Minnesota, Washington University in St. Louis, and University of Colorado-Boulder sites, which all use 3T Siemens MRI scanners. We excluded one twin site as it uses a different platform with different scanner parameters.

Despite evidence of decent cross-platform reliability for structural and functional imaging (Duchesne et al., 2019; Schwartz et al., 2019; Keenan et al., 2021), we chose to harmonize our analyses on data collected from a single platform type (Casey et al., 2018). ABCD study-wide exclusion criteria include a diagnosis of schizophrenia, moderate to severe autism spectrum disorder, intellectual disabilities, major and persistent neurological disorders, multiple sclerosis, sickle cell disease, seizure disorders like Lennox-Gastaut Syndrome, Dravet Syndrome, and Landau Kleffner Syndrome, or substance abuse disorders at time of recruitment. Subjects with mild autism spectrum diagnosis, history of epilepsy, traumatic brain injury, and MR incidental findings were excluded from the present analysis.

### Resting state fMRI data collection

Structural and functional data from the three included sites were acquired using Siemens Prisma 3T scanners with a 32-channel head coil. Detailed acquisition parameters are previously described in the literature (Casey et al., 2018; Hagler et al., 2019). Scan sessions consisted of a high-resolution (1 mm^3^) T1-weighted image, diffusion weighted images, T2-weighted spin echo images (1 mm^3^), resting-state fMRI (rs-fMRI), and task-based fMRI. We utilized rs-fMRI solely in this study. rs–fMRI data were collected with an echo-planar imaging sequence with 2.4 mm voxels, TR = 800 ms, TE = 30 ms, and multiband slice acceleration factor = 6. Participants completed up to four runs of 5-minute resting state scans. Framewise integrated real-time MRI monitoring (FIRMM (Dosenbach et al., 2017)) was used to monitor subject head motion during data collection, and scan operators may have stopped resting-state data collection after three runs if 12.5 minutes of low-motion resting-state data had been collected. Thus, variable numbers of rs-fMRI runs were collected per subject.

Additional exclusions were made based on poor structural scan quality (determined with curated data release 2.0.1 sheet *freesqc01.txt*) or low quality/high motion resting state fMRI (determined as a score of zero for *fsqc_qc* or greater than one for *fsqc_qu_motion*, *fsqc_qu_pialover*, *fsqc_qu_wmunder*, or *fsqc_qu_inhomogeneity*), as reported in prior studies (Rosenberg et al., 2020). Participants with fewer than 900 brain volumes of rs-fMRI data passing these measures for FreeSurfer reconstruction were also excluded. ABCD Data Release 2.0.1 used FreeSurfer version 5.3 for quality control checks (Fischl, 2012). After filtering, we used resting state fMRI and behavioral data from 1,115 unique subjects (552 male), including 170 pairs of monozygotic (MZ) twins, 219 pairs of dizygotic (DZ) twins, and 337 singletons. All twin pairs were same-sex, and all subjects were 9–10 years old. Included singletons were also balanced for ancestry, ethnicity, sex, and pubertal development with twins (Table 1).

**Table 1:** Participant information and cohort breakdown. Data were acquired from the publicly available ABCD data release 2.0.1. Participants were divided into cohorts to balance unrelated participants with the demographics of the twin participants, while respecting the demographic diversity of individual ABCD sites. Percent ancestry values reflect the percentage of a participant’s genetic makeup attributed to a given ancestry, then averaged across participants. Ethnicity values represent a 4-level race/ethnicity model, where participants self-identified as White, Black, Hispanic, Asian, or other, and these coded values were averaged across participants.

|  | Total included | Monozygotic pairs | Dizygotic pairs | Unrelated test | Unrelated train |
| --- | --- | --- | --- | --- | --- |
| Total | 1,115 | 170 | 219 | 526 | 200 |
| Site 20 | 387 | 75 | 86 | 184 | 60 |
| Site 21 | 371 | 34 | 81 | 166 | 90 |
| Site 22 | 357 | 75 | 68 | 176 | 50 |
| Age (mos, *interview_age*) | 121.6 | 122.5 | 122.0 | 121.6 | 120.7 |
| % female (*gender*) | 50 | 52 | 50 | 50 | 50 |
| % Ancestry: African (*AFR*) | 12.7 | 9.2 | 11.8 | 11.3 | 20.4 |
| % Ancestry: European (*EUR*) | 80.5 | 82.5 | 82.6 | 81.4 | 73.3 |
| % Ancestry: American (*AMR*) | 4.9 | 6.7 | 4.0 | 5.2 | 3.8 |
| % Ancestry: East Asian (*EAS*) | 2.0 | 1.6 | 1.6 | 2.1 | 2.5 |
| Ethnicity (*race_ethnicity*) | 1.667 | 1.658 | 1.63 | 1.73 | 1.58 |

### Preprocessing

rs-fMRI data were downloaded via the DCAN Labs ABCD-BIDS Community Collection (Feczko et al., 2021) (NDA Collection 3165). This is a regularly updated dataset of ABCD Brain Imaging Data Structure (BIDS) (Gorgolewski et al., 2016) version 1.2.0 pipeline inputs and derivatives, using source data from the ABCD Study participants baseline year 1 arm 1 DICOM imaging data that passed initial acquisition quality control from the ABCD Data Analysis and Informatics Center (DAIC) (Hagler et al., 2019) and retrieved from the NIMH Data Archive (NDA) share of ABCD fast-track data (NDA Collection 2573). DICOMs were converted to BIDS input data using Dcm2Bids (Bedetti, 2022), which reorganizes NiftiImages produced with dcm2niix (Li et al., 2016). Raw images were preprocessed using the DCAN Labs ABCD-BIDS MRI processing pipeline (Sturgeon et al., 2021) (for details, see github.com/DCAN-Labs/abcd-hcp-pipeline; osf.io/89pyd), which is based on the Human Connectome Project (HCP) Minimal Preprocessing Pipeline (Glasser et al., 2013) with additional modifications specifically for the ABCD Study dataset and summarized below.

The first stage of the pipeline, PreFreeSurfer, performed brain extraction, alignment, and N4 bias field correction on the T1w and T2w images. The second stage, FreeSurfer (Dale et al., 1999; Fischl, 2012), segmented the resulting T1w images and identified tissue boundaries to register to a FreeSurfer template and produce brain masks. The third stage, PostFreeSurfer, used the brain masks to register T1w images to MNI space using ANTs symmetric image normalization method (Avants et al., 2008). Surfaces were then transformed to standard space using spherical registration and converted to CIFTI format along with the standardized brain volumes. Multimodal surface registration was performed with MSM-sulc, the state-of-the-art surface-based alignment procedure (Robinson et al., 2014). By using MSM-sulc, we expect regional boundaries to be well-aligned across brains, thus minimizing the potential influence of anatomical differences on differences in functional connectivity (Coalson et al., 2018). The fMRIVolume stage performed functional image distortion correction using reverse phase-encoded spin echo images to correct for local field inhomogeneities. Eta squared values were computed for each image to a participant-level average of all field maps, and the pair with the highest value (i.e., most representative of the average) was selected to avoid potential motion confounds. Finally, the fMRISurface stage performed 2-mm full-width half-max spatial smoothing. DCAN BOLD Processing (DCAN_Labs, 2022) was used to perform standard data processing.

First, fMRI data were demeaned and detrended with respect to time. Covariates of no interest were regressed from the data, including mean white matter, cerebrospinal fluid, overall global signal, mean grayordinate timeseries, and 6-movement variables (X,Y,Z translation and roll, pitch, yaw rotation). Global signal regression has been shown to strengthen associations between RSFC and behavioral variables (Li et al., 2019). Timeseries data were then band-pass filtered between 0.008 and 0.09 Hz using a 2nd order Butterworth filter. Data were then filtered for respiratory motion for the frequencies (18.582 to 25.726 breaths per minute) of the respiratory signal, which has been shown to improve the stability of framewise displacement (FD) estimates (Fair et al., 2020). “Bad” frames, where motion exceeds FD of 0.3 mm, were removed when demeaning and detrending such that denoising betas were only calculated for “good” frames. For the band-pass filtering, interpolation was used to replace those frames, so that preprocessing of the timeseries only includes “good” data but avoids aliasing due to missing timepoints. Finally, the filtered timeseries were normalized within run and concatenated across runs. The derivatives used in this analysis are *ses-baselineYear1Arm1_task-rest_bold_desc-filtered_timeseries.dtseries.nii. Participant inclusion pipeline

### Connectivity hyperalignment

We used connectivity hyperalignment (CHA) (Guntupalli et al., 2018) to functionally-align fine-scale connectivity information across subjects. Given the small quantity (1480 *±* 142 brain volumes; *mean* ± *s*.*d*.) of resting state fMRI data available per subject, we chose to train the hyperalignment model common space using a cohort of 200 subjects who were then excluded from further analysis. These 200 training subjects were singletons enrolled at one of the three included sites and were matched with twin subjects on gender, pubertal development, age, and race/ethnicity (see Table 1 and Figure 2). The remaining 915 subjects (including 137 singletons and the 389 twin pairs) were used for subsequent analyses (“test” subjects). Crucially, the hyperalignment model space never saw data from any test subjects during training.

**Figure 1:**
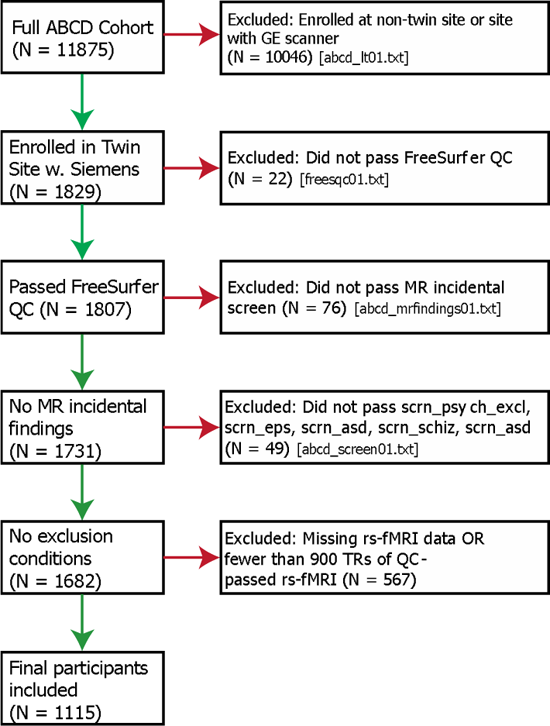
Flowchart illustrating participant inclusion/exclusion criteria.

**Figure 2:**
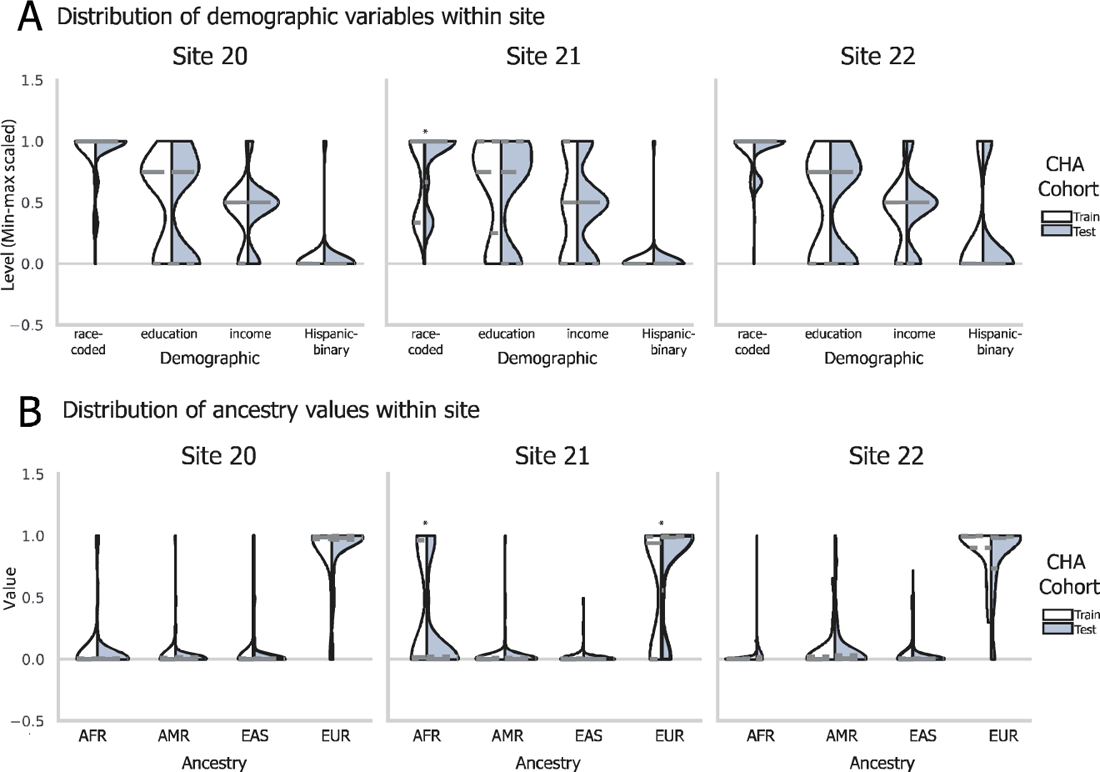
Balancing of participant demographics and genetic ancestry across hyperalignment training. The three sites differed in mean demographic attributes, including a 4-level race model, parental education level, parental income, and Hispanic identity (A) and in mean genetic ancestry (B). To respect this diversity and sample the entire range of demographics in the normative hyperalignment model, we selected subjects to train the CHA model to reflect sample demographics along these four dimensions.

CHA aligns fine-scale connectivity patterns in a cortical field to the same targets elsewhere in the brain. While computing connectivity profiles to be hyperaligned, it is often desirable to use individualized connectivity targets to account for topographic idiosyncrasies in target regions. Individualized connectivity targets can be generated with individualized parcellations (Glasser et al., 2016; Langen et al., 2018; Kong et al., 2019, 2021, 2023; Anderson et al., 2021) or by iterating the hyperalignment algorithm (Busch et al., 2021; Jiahui et al., 2023). In our implementation for this study, we used parcels from the Glasser cortical parcellation (Glasser et al., 2016) as the cortical fields and targets to be hyperaligned. The Glasser parcellation provides whole-cortex coverage and has been used in prior studies evaluating RSFC and behavioral predictions (Dubois et al., 2018a, 2018b; Feilong et al., 2021). We co-registered subjects’ brains using surface-based alignment with the MSM algorithm (Robinson et al., 2014) which aligns the region boundaries well (Coalson et al., 2018), but future work may want to use individualized connectivity targets instead of parcels, particularly if using vertex-level connectivity targets (Feilong et al., 2021).

We trained a hyperalignment model for each parcel using the 200 training subjects’ “fine scale” connectomes, which capture the connectivity between connectivity seeds (each vertex in the cortical field defined by the parcellation) with connectivity targets across the rest of the brain (the average timeseries of each other parcel) (Figure 3). For example, for a parcel with 430 vertices, its fine-scale connectivity matrix would be a matrix of dimensions 430 by 359 correlation coefficients, where 430 corresponds to the number of vertices within the parcel and 359 corresponds with each other region in the parcellation. The same parcel’s coarse-scale connectivity profile would be a vector of 359 correlation coefficients (Figure 3B). During training of the hyperalignment space, the 359 pattern vectors representing the connectivity targets of one brain are hyperaligned to another brain’s vectors for the same field and targets with a high-dimensional transformation that minimizes the distances between vectors to the same targets. We calculate these transformations using the Procrustes transformation to derive the optimal high-dimensional improper rotation that minimizes these distances, preserving their geometry. A template connectivity information space for a cortical field is calculated for a normative sample of brains by first hyperaligning brain 2 to brain 1, then hyperaligning brain 3 to the mean patterns for hyperaligned brains 1 and 2, and so forth. In a second iteration, each brain is hyperaligned to the mean connectivity pattern vectors from the first iteration. The mean pattern vectors for each cortical field after the second iteration serves as the template for hyperaligning other brains to this high-dimensional model connectivity space (“Common model space for parcel *p*” in Figure 3A). A new individual-specific transformation is calculated for each new brain (*R* for each test subject in Figure 3A). Each vertex in this model template serves as a model dimension, and the individual-specific transformation resamples the vertices in a new participant’s brain into these model dimensions as weighted averages.

**Figure 3:**
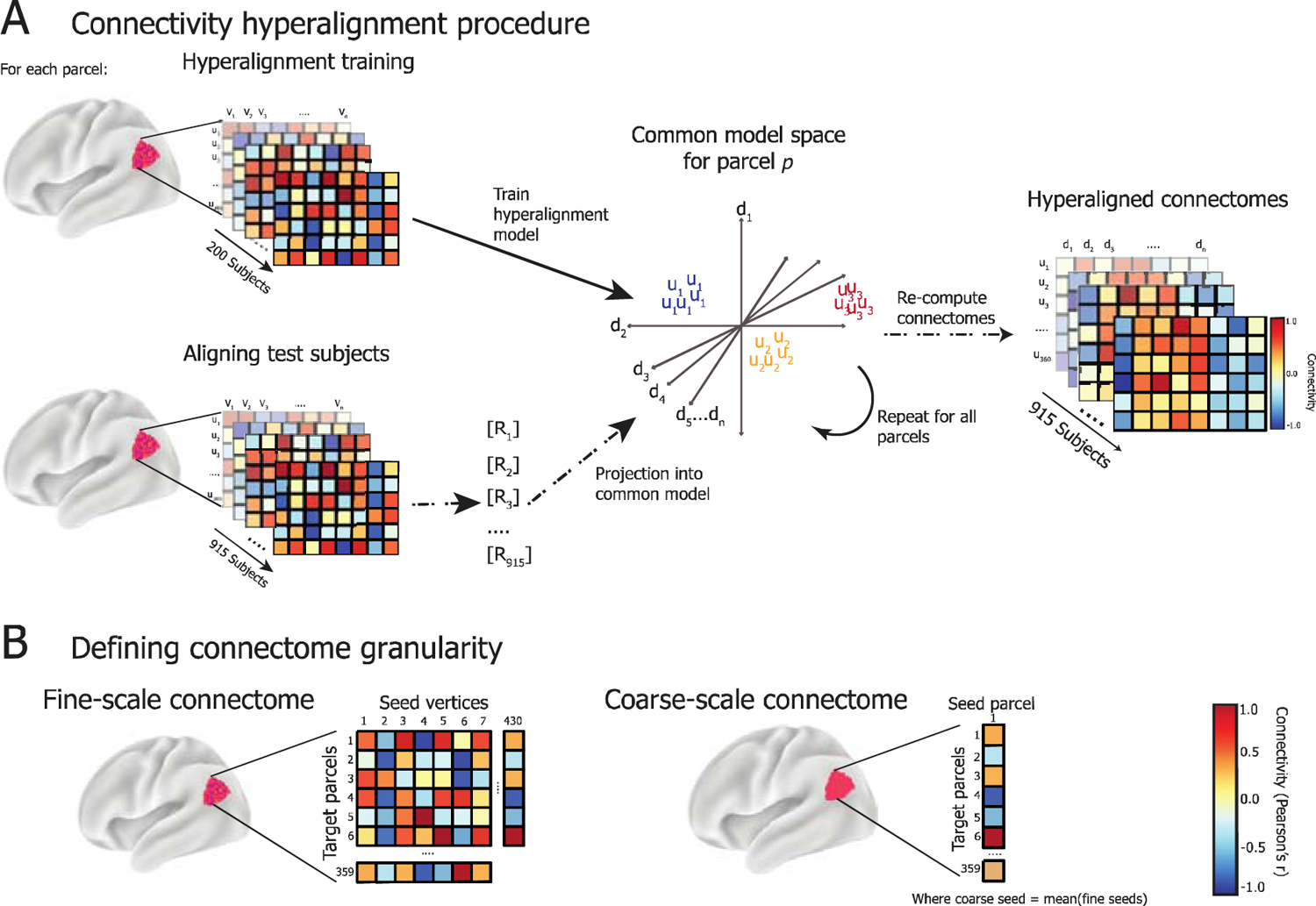
A. Connectivity hyperalignment training procedure. For each parcel in the Glasser parcellation (here, region 329 with 430 cortical vertices), connectivity matrices are computed as the strength of connectivity (Pearson’s correlation) between the activity timeseries of vertices within the parcel (seeds; *u*) and the average activation across vertices in all other parcels (targets; *v*). The connectivity matrices for 200 held-out, unrelated subjects are used to train the hyperalignment common model space for a given parcel. The model space is of dimensions *D* = [*d*_1_…*d*_n_] which are weighted combinations of *U* = [*u*_1_…*u*_n_], the dimensions of each subject’s connectivity matrices in their native anatomical space. *D* represents the space where connectivity targets are best aligned across training subjects. Then, connectomes for the 915 test subjects are projected into *D* via subject-specific transformation matrices *R*, which detail a mapping from given subject *i*’s anatomical space *U*_i_ into *D*. This procedure is repeated to derive *R* for each test subject and parcel. Then, the regional activation pattern for each parcel in anatomical space is mapped via *R* into the region’s common space to compute the hyperaligned timeseries for each subject, which is then used to recompute connectivity matrices between hyperaligned seeds *v* and model dimensions *d*. B. Connectome granularity. Fine-scale connectomes are multivariate patterns of the connectivity weights between the time-series responses of all individual vertices in a given parcel and the average time-series response of each other parcel in the brain. Coarse-scale connectomes are vectors (univariate), representing the connectivity weight between the average time-series response of a given parcel, averaged across all of its vertices, and those of each other parcel in the brain.

For each region, we trained a separate high-dimensional common model space *D* based on the fine-scale functional connectomes of the 200 training subjects. The dimensionality of the model space is equal to the number of vertices, but the model dimensions correspond to shared functional connectivity properties across individuals instead of anatomical locations (Haxby et al., 2020). After the model dimensions were learned, we discarded training subjects’ data from downstream analyses. We derived invertible transformation matrices *R* for the testing subjects’ connectomes into the model space, which map vertex timeseries from the anatomically-defined, vertex dimensions into the functionally-defined, model dimensions. The derivation of *R* for testing subjects was determined based on the analysis we performed to ensure that *R* was not overfit and outlined in the methods section for each analysis.

Shown here are the distributions of participants’ scores, scaled for visualization, for the participants allocated for CHA train or test cohorts. Violins show training cohort on left and testing cohort on right, with quartiles of each distribution shown with a dashed line. Significance of differences along each dimension between CHA cohorts were tested with a two-tailed independent sample t-test and * denotes *p <* 0.05 after Bonferroni correction.

### Functional connectomes

After aligning data from test subjects into the CHA common space, we re-computed coarse and fine-scale functional connectivity matrices, where the fine-scale patterns capture model dimensions instead of vertices for each parcel. We defined fine-scale functional connectivity profiles as in the input data for the CHA training procedure: the correlation between the timeseries of all 59,412 model dimensions with the average timeseries of each parcel. This results in a matrix of 59,412 by 360 connectivity weights per subject, and we then sort the 59,412 dimensions into their parcels to get a parcel’s fine-scale connectome after CHA (Figure 3B).

We defined coarse-scale functional connectivity profiles as the correlation between the average timeseries for all pairs of parcels, resulting in a matrix of 360 by 360 correlation coefficients per subject (Figure 3B). Before hyperalignment, this averaging is performed over the vertices within a parcel. After hyperalignment, this averaging is performed over the model dimensions within a parcel. The fine-scale and coarse-scale connectomes were computed on data pre- and post-hyperalignment (subsequently referred to as anatomical alignment [AA] and connectivity hyperalignment, respectively) for each subject to evaluate the effect of functional alignment and connectome granularity on heritability, reliability, and behavioral prediction.

### Individual difference matrix (IDM) reliability

Developmental neuroimaging data is associated with high levels of noise and low reliability (Power et al., 2012; Satterthwaite et al., 2012; Birn et al., 2013; Fair et al., 2020; Kennedy et al., 2022), as are smaller (vertex-resolution) connectivity seeds (Haak et al., 2018). Variability in RSFC can reflect idiosyncrasies driven by cortical communication patterns indicative of neurocognitive variability, but it can also reflect artifacts due to data acquisition noise, movement (Power et al., 2012; Van Dijk et al., 2012), and functional–anatomical mismatch. In adult populations, brain parcellations (e.g., Glasser parcellation (Glasser et al., 2016)) average adjacent areas of cortex to improve signal-to-noise and reduce the dimensionality of the data, and adult-defined parcellations are frequently applied to development datasets (Chen et al., 2022; Marek et al., 2022). This approach assumes functional–anatomical correspondence will improve at the group level and that functional boundaries are constant across development, and comes at the cost of lower granularity.

Another approach to group alignment, hyperalignment has been shown to improve the reliability of fine-scale information and individual differences by correcting more thoroughly for anatomical variability through a space that aligns patterns of functional connectivity in high-dimensions (Dubois and Adolphs, 2016; Haxby et al., 2020). To our knowledge, hyperalignment or other functional alignment approaches have yet to be applied to developmental fMRI data, and we hypothesized it could yield access to meaningful individual differences in RSFC patterns. We measured the effect of CHA on reliability of individual differences using an individual difference matrix (IDM) reliability analysis.

We calculated the reliability of individual differences in functional connectomes as follows. For the 526 unrelated test subjects, we computed fine-scale functional connectomes by splitting each subject’s RS timeseries in half to create two “pseudo-sessions” (mean ± std. across subjects = 742 *±* 71 TRs), resulting in two fine-scale functional connectomes for each subject. These initial matrices served as the fine-scale AA functional connectomes, and averaging over the vertex dimensions yielded two coarse-scale AA connectomes.

To perform CHA on a given test subject’s data, we took their two pseudo-session fine-scale AA connectomes and derived two transformation matrices *R* independently for each connectome. These *R* matrices were then applied to their respective pseudo-session timeseries data to rotate the subject’s resting-state timeseries into the common dimensions. This was repeated for all parcels, keeping each pseudo-session entirely independent, before recomputing fine-scale and coarse-scale functional connectomes for each pseudo-session. After this procedure, each subject has 8 different connectomes per parcel: AA fine, AA coarse, CHA fine, and CHA coarse, which were independently processed for each half of their timeseries data.

Then, for each connectome and parcel, we vectorized the functional connectomes and computed individual difference matrices (IDMs) for each pseudo-session. IDMs are subject-by-subject pairwise dissimilarity matrices, where each value in the matrix is the correlation distance (1 - Pearson’s *r*) between two subjects’ regional connectivity profiles. After computing IDMs, we took the upper triangle of the IDM and correlated them (Pearson’s *r*) across split halves (but within connectome type) to measure the reliability of the individual differences in the functional connectomes across the splits (Feilong et al., 2018). This metric reflects the reliability of the *idiosyncrasies* in a subject’s functional connectomes, where subjects are more similar to themselves across split halves than they are to any other subject, rather than a standard reliability analysis. We used a Mantel test (Mantel, 1967) implemented in Python (https://github.com/jwcarr/mantel) to assess statistical significance of IDM reliability using 10,000 permutations. This analysis results in four spatial maps of IDM reliability scores, with one score for each parcel within the AA fine, AA coarse, CHA fine, and CHA coarse connectome types.

### Multidimensional heritability analysis

In this analysis, we assessed the heritability of functional connectomes using RS data from 389 twin pairs, including 170 MZ pairs and 219 DZ pairs. RSFC heritability for each parcel was estimated for the AA fine, AA coarse, CHA fine, and CHA coarse-scale connectomes, as in the prior reliability analysis, to assess the degree of genetic control over the information represented by each connectome type. CHA was performed using the same model space as the reliability analysis. The 778 subjects’ data included in the heritability analysis were hyperaligned using their entire timeseries, rather than the pseudo-sessions used to assess reliability.

Functional connectomes at vertex-granularity are an inherently high-dimensional phenotype; when dimensionality of the phenotype is collapsed, the reliability of individual differences decreases (Feilong et al., 2018). To offer greater statistical power in analyzing the heritability of this high-dimensional phenotype, we used a multidimensional estimate of heritability (Ge et al., 2017; Anderson et al., 2021) to summarize the degree to which functional connectivity profiles for each parcel are under genetic control. This model takes a phenotypic similarity matrix, which is computed by vectorizing the connectomes for each parcel and computing the pairwise correlation of connectomes across all 778 subjects. It also takes a genetic kinship matrix, which we estimated with SOLAR Kinship2 (Almasy and Blangero, 1998), to estimate h^2^-multi, or the score indicating heritable control over RSFC. This was computed for each parcel at each scale and alignment.

### Neurocognition PC prediction analysis

Prior work has shown that, after hyperalignment, fine-scale connectomes better predict individual differences in cognitive ability than coarse-scale connectomes in adults (Feilong et al., 2021). We asked whether this fine-scale information also reflects information more predictive of cognition in children, where the fine-scale connectome is likely more obscured by collection noise yet sculpted with experience. Further, we asked whether there is a relationship between the type of cognition being predicted and the connectome scales that best capture variation in that phenotype.

Cognitive abilities are among the most heritable dimensions of human behavior (Plomin and Deary, 2015). Heritability (h^2^) of cognitive abilities varies by domain and increases with age, with higher estimates for general cognitive ability (h^2^ = 0.54–0.85 at 10–12 years (Bouchard, 2004) and h^2^ = 0.60–0.73 at 8–21 years (Mollon et al., 2021)) than for specific cognitive abilities like learning/memory (h^2^ = 0.18–0.55 at 8–21 years (Plomin and Spinath, 2004) and h^2^ = 0.39 at 18–67 years (Fletcher et al., 2014)), which tend to be more sensitive to experiential factors. Thus, we looked at whether the variation in reported heritability between general cognitive ability and learning/memory relates to the functional significance of our heritability and reliability findings.

We operationalized general cognitive ability and learning/memory using the neurocognition principal components (neurocog PCs) identified by Thompson et al., (2019) and made available through the ABCD Study data repository. Thompson et al. (2019) identified composite principal components of neurocognition using Bayesian Probabilistic Principal Components Analysis (BPPCA) over the NIH Cognitive Toolbox tasks administered in the ABCD study, which measure episodic memory, executive function, attention, working memory, processing speed, and language abilities. The BPPCA yielded a three-component solution, roughly corresponding to general cognitive ability, executive function, and learning/memory. Since the NIH Toolbox tasks that canonically measure executive function did not load onto the executive function component in this solution, we excluded this component from our analyses, instead focusing on general cognitive ability and learning/memory. Tasks loading onto the general cognitive ability component include Toolbox Picture Vocabulary, Toolbox Oral Reading test, List Sort Working Memory task, and Little Man task. Tasks loading onto the learning/memory component include the Toolbox Picture Sequence Memory task, the RAVLT total number correct, and the List Sort Working Memory task (Thompson et al., 2019). Heritability of composite scores were estimated with Falconer’s formula (Falconer, 1996) and with SOLAR Polygenic (Almasy and Blangero, 1998) testing sex, age, ethnicity, and site as covariates.

We predicted individual subjects’ scores onto the neurocognition principal components based on individual differences in RSFC from our cohort of 526 unrelated subjects using a principal component ridge regression adapted from our previous study (Feilong et al., 2021). For each parcel, scale, and alignment, we computed subject-wise individual differences matrices (IDMs) using functional connectivity over the entire resting-state timeseries, which captures the covariance structure among subjects’ functional connectomes. Next, we ran a principal components analysis (PCA) over the IDM to decompose the IDM into main orthogonal dimensions along which individuals’ connectivity profiles differ from one another. We then trained a ridge regression model to predict neurocognition PC loadings from the decomposed IDM. Models were trained and tested using nested cross-validation, where the inner fold optimized regression model parameters (including the number of IDM PCs retained and α) across three sub-folds and the outer fold scored the selected model on a held-out subject. Candidate regression model parameters were distributed evenly on a logarithmic scale, from 10 to 320 PCs and no PCs (no dimensionality reduction), and α chosen from 121 values between 10*^−^*^20^ to 10^40^. Models were evaluated using the cross-validated coefficient of determination *R*^2^, which shows the variance in neurocognition PC loadings accounted for by the prediction models. To assess statistical significance of model performance against chance, we used permutation testing with 1,000 iterations of shuffling the neurocognitive scores, performed at the parcel level. To compare prediction scores across connectome types, we generated a null distribution of the difference in scores between pairs of connectome types by randomly permuting the connectome label 1,000 times and recomputing the mean difference, then calculating a two-tailed p*-*value of the true difference relative to the null distribution. Between-connectome p*-*values were corrected for multiple comparisons using the Bonferroni method.

Prior work has indicated that in-scanner head movement is correlated with neurocognition scores. We implemented strict participant exclusion based on data quality control, but still found a mild relationship between mean framewise displacement (FD) and neurocognitive scores. To control for possible effects of movement on our prediction, we repeated the neurocognition PC prediction analysis by including FD as a covariate when comparing observed and predicted scores.

### Statistical tests

Within each analysis, we estimated the statistical significance of the score calculated for each parcel and connectome type. For the reliability analysis, we calculated the statistical significance of the correlation between IDMs using a Mantel test (Mantel, 1967) with 10,000 permutations and implemented in Python (https://github.com/jwcarr/mantel). For the heritability analysis, we calculated statistical significance of h^2^-multi scores using subject-based permutations, where the kinship matrix was randomly shuffled 1,000 times (Anderson et al., 2021). For the prediction analysis, we calculated statistical significance of *R*^2^ values by permuting the neurocognitive scores 1,000 times. All p-values were then false discovery rate adjusted and surface plots are all thresholded at *q <* 0.05.

In each of the reliability, heritability, and prediction analyses, we compute p-values to compare the scores at each parcel across connectome type. We did this by generating a null distribution of the difference in scores between pairs of connectome types by randomly permuting the connectome label 1,000 times and recomputing the mean difference of the scores. We then compared the true difference between scores relative to the null distribution and computed a two-tailed p-value. This was repeated for each pair of connectome types and p-values, as reported, were corrected for the six multiple comparisons using the Bonferroni method.

### Data availability

The ABCD study is longitudinal, so the data repository changes over time and can be found here: nda.nih.gov/abcd. In the current study, we used the ABCD data release 2.0.1, downloaded via ABCD-BIDS Community Collection (Feczko et al., 2021) NDA Collection 3165.

### Code availability

ABCD Study data preprocessing code can be found here: github.com/DCAN-Labs/abcd-hcp-pipeline. ABCD Study data processing code can be found here: github.com/DCAN-Labs/dcan_bold_processing. Code for running the multiscale heritability analysis can be found here: github.com/kevmanderson/h2_multi. Code for running the nested PCA ridge regression was based off: github.com/feilong/IDM_pred. Hyperalignment was performed with PyMVPA: www.pymvpa.org. All analysis code specific to this study will be released upon publication.

## Results

### Establishing the reliability of individual differences

Here, we analyzed RSFC in a large cohort of 1,115 children, aged nine to ten, using two methods of group alignment. The first uses a standard approach to align data in a common space based on anatomical location (anatomical alignment [AA]), while the second approach uses connectivity hyperalignment (CHA) (Guntupalli et al., 2018) to learn a common information space based on fine-scale patterns of functional connectivity within cortical fields (Glasser parcels) via an adapted Generalized Procrustes Analysis (see Methods for details). CHA allows for local remixing of vertex-wise functional connectivity profiles (vertices’ vectors of connectivity to targets) within an anatomically constrained region (Haxby et al., 2011), thereby projecting idiosyncratic patterns of vertex connectivities to the same target into a common connectome space (Haxby et al., 2020). In contrast with coarse alignment, which averages time-series responses across a region’s vertices to calculate a single connectivity strength with each target region, CHA retains the vertex-by-vertex variation on connectivity strength to each target region (Figure 3B). Within this common, fine-scale space, individual differences in functional connectivity become more reliable (Feilong et al., 2018) and more predictive of cognitive traits (Feilong et al., 2021) in adults.

To address the question of individual differences in fine-scale functional connectivity among children, we use CHA to build a model of shared brain function. This negates concerns about differences in cortical development or anatomy that may confound modeling of functional connectivity patterns, and accounts for the noise and other undesirable idiosyncrasies endemic in developmental neuroimaging. To do this, we used CHA to build a template common connectome for each parcel on a training sample of 200 unrelated children, counterbalanced across included sites and representative of demographics within site (Table 1; Figure 2). We then hyperaligned each of the 915 remaining connectomes to this common connectome, using either entire or split-half rs-fMRI timeseries (depending upon the downstream analysis). Note that this common information space is based on children of the same age and demographics as the test sample, and that the test sample’s data played no role in deriving the common space. We addressed the relative effects of hyperalignment versus simply higher-dimensional measurements by averaging the functionally aligned CHA data and by retaining the high-dimensional, anatomically aligned data, to have a full comparison of connectomes that are aligned anatomically, at both the coarse-scale and fine-scale level, and connectomes that are CHA-aligned, at both the coarse-scale and fine-scale level.

We first assessed whether idiosyncrasies in cortical activity patterns were more reliable after fine-scale hyperalignment, as has been shown in adult cohorts (Feilong et al., 2018). Here, we define idiosyncrasies in terms of individual differences matrices (IDMs), or the pairwise correlation distance between all pairs of subjects. IDMs capture the dissimilarity structure (or idiosyncrasies) of functional connectivity patterns across subjects. We assessed the reliability of these idiosyncrasies by splitting each subject’s RS timeseries in half to create two “pseudo-sessions” and calculating independent connectomes for each split-half. For hyperalignment, we aligned each subject’s two pseudo-session connectomes to the common space independently, treating them as we would two separate subjects, as to not bias the reliability estimates. We then computed an IDM for each pseudo-session and correlated the two pseudo-sessions IDMs to measure the reliability of individual differences. This process was performed for anatomically aligned (AA) coarse- and fine-scale connectomes and for CHA coarse- and fine-scale connectomes separately.

CHA, particularly at the fine scale, improved the reliability of individual differences in functional connectivity over anatomical alignment alone (Figure 4). Across all parcels, reliability of coarse connectomes increased from an average of *r* = 0.34 *±* 0.1 for AA to *r* = 0.58 *±* 0.08 for CHA (*p <* 0.0001). Reliability of fine connectomes increased from an average of *r* = 0.43 *±* 0.11 for AA to *r* = 0.86 *±* 0.07 for CHA. After CHA, 100% of parcels at both scales showed greater reliability, and 99% of parcels showed greater reliability at the fine scale than the coarse scale. Notably, all four types of connectomes showed high split-half reliability, with CHA affording the greatest reliability. This result establishes that hyperalignment affords access to reliable information within an individual child’s fine-scale functional connectome, affording confidence in subsequent results based on these connectomes.

**Figure 4:**
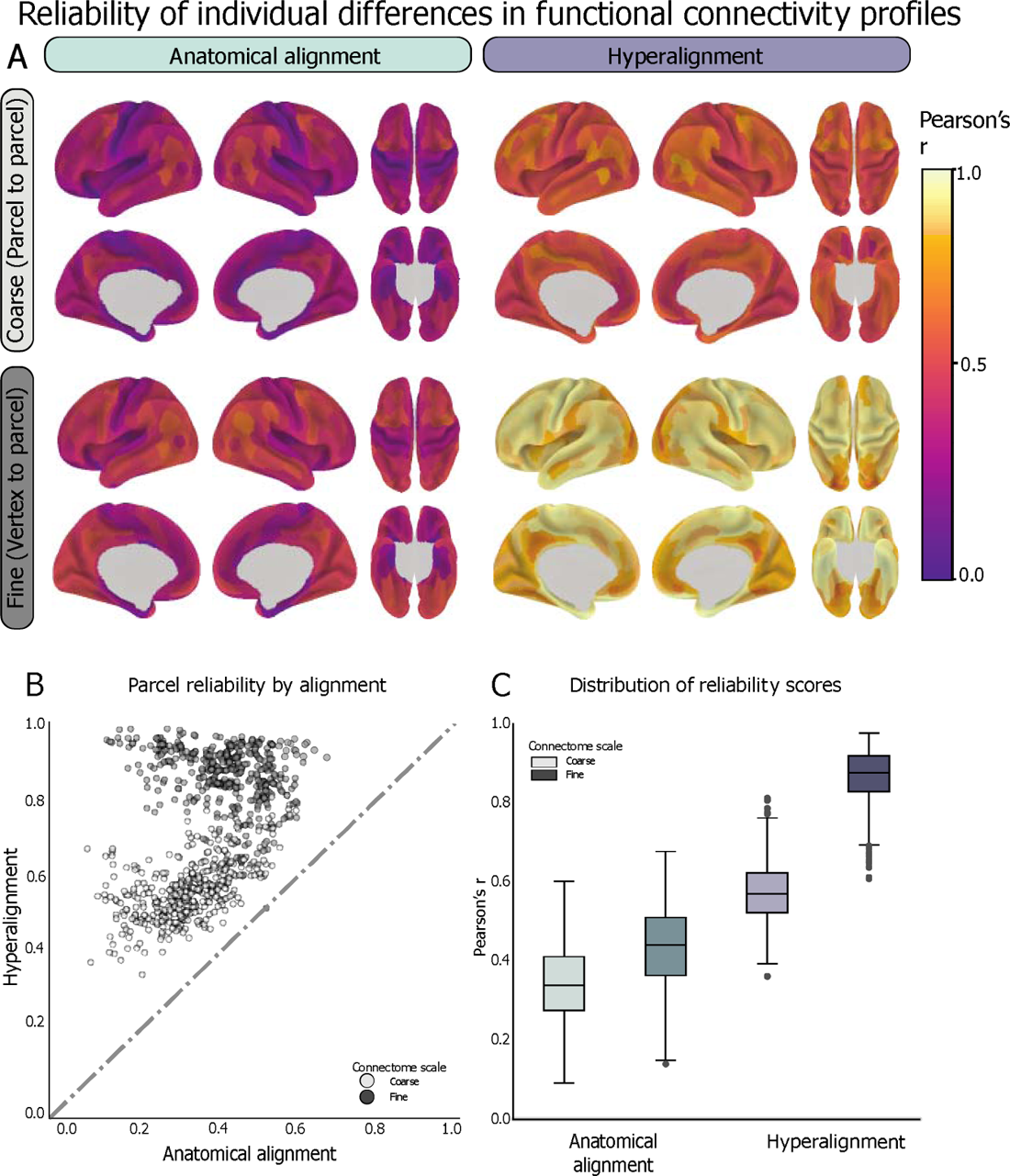
The reliability of individual differences in functional connectivity profiles was analyzed with data from 526 unrelated subjects. Subjects’ timeseries data for each parcel were split into two pseudo-sessions, which were hyperaligned into the trained common space independently. For each region, split-half timeseries data pre- and post-hyperalignment were used to compute functional connectivity profiles at the fine and coarse scales. Individual differences matrices (IDMs) were computed across all pairs of subjects for each data split. IDMs were then correlated (Pearson’s *r*) across splits and repeated for each parcel. This process was repeated for all parcels at the coarse and fine scales, before and after hyperalignment. A. Spatial distribution of reliability scores before and after hyperalignment at the fine and coarse scales, as visualized on the cortical surface for each parcel. B. The reliability of individual differences increases after hyperalignment for all parcels. 99% of fine connectivity profiles (darker) are more reliable than coarse ones (lighter) regardless of alignment type. C. Box-and-whisker plots showing the distribution of IDM reliability scores across parcels within connectome type. Box shows first quartile, median, and third quartile of distribution. Whiskers represent 1.5 times the interquartile range, and points represent outlier values.

### Assessing heritability of connectome granularity

We asked whether the more reliable fine-scale connectome simply reflected the same information about cortical functional architecture at higher magnification or a different class of information that may be more or less heritable. We investigated this question by calculating the heritability of the two coarse-scale and the two fine-scale connectomes. We hypothesized that heritable factors determine the skeletal attributes of the connectome and are more evident at the coarse spatial scale (Lenroot et al., 2009; Blokland et al., 2012; Schmitt et al., 2014), whereas the detailed configurations of connectomes reflect the interaction of the heritable connectome with unique experiential factors producing variability at a finer spatial scale.

The heritability analysis used RSFC data from 778 subjects comprising 219 dizygotic twin pairs and 170 monozygotic twin pairs (Table 1). As in prior analyses, the connectivity hyperalignment model was trained on 200 unrelated subjects, then the connectomes of the 778 twin subjects were hyperaligned to this model. Functional connectomes of vertex-granularity are an inherently high-dimensional phenotype; thus, to explicitly model the heritability of this phenotype, we used a multidimensional estimate of heritability (h^2^-multi) (Anderson et al., 2021) to model both the phenotypic similarity matrix between participants as well as the genetic kinship matrix (see *Methods: Multidimensional heritability estimate* for more information).

We find that both the AA and CHA coarse-scale connectomes were more heritable than the more reliable fine-scale connectomes (mean coarse-scale h^2^-multi = 0.18 *±* 0.05, mean fine-scale h^2^-multi = 0.10 *±* 0.03, *p <* 0.0001), with 99 % of parcels showing greater heritability for coarse-scale connectomes (Figure 5). At the coarse scale, anatomically aligned functional connectomes are significantly more heritable than hyperaligned connectomes (AA h^2^-multi = 0.20 *±* 0.05; CHA h^2^-multi = 0.16 *±* 0.04; *p <* 0.0001). These results indicate that connectivity information aligned coarsely based on anatomical features are under strong genetic control, consistent with previous studies showing genetic control over cortical anatomy in children (Peper et al., 2009) and network-level RSFC (Glahn et al., 2010).

**Figure 5:**
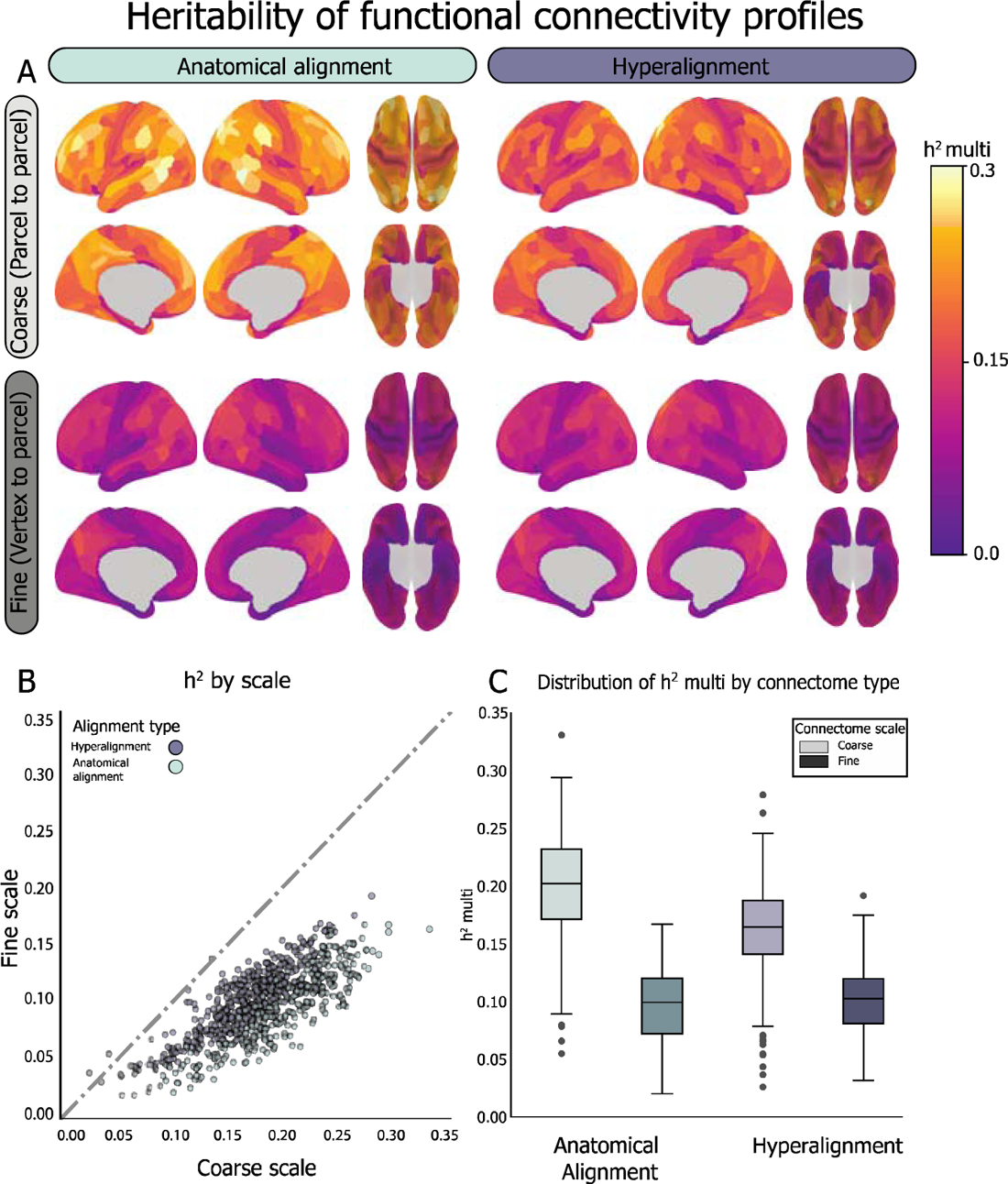
Heritability of functional connectivity was estimated using a multidimensional heritability model (Anderson et al., 2021) using resting-state fMRI data from 778 9–10-year-olds (170 MZ pairs and 219 DZ pairs). Functional connectivity between a given parcel and all other parcels in the brain was calculated at fine and coarse granularities, before and after connectivity hyperalignment. The coarse-scale connectivity profiles aligned based on cortical anatomy are the most heritable. A. Spatial distribution of h^2^-multi scores before and after hyperalignment at the fine and coarse scales, as visualized on the cortical surface for each parcel. B. For both anatomically aligned and hyperaligned data, connectivity profiles are more heritable when calculated at the coarse scale than the fine scale (99.6 % of parcels). C. At the coarse scale, the anatomically aligned connectivity profiles are more heritable than the hyperaligned ones, but not at the fine scale, which is under comparatively minuscule genetic control. Box-and-whisker plots showing the distribution of h^2^-multi across parcels within connectome type. Box shows first quartile, median, and third quartile of distribution. Whiskers represent 1.5 times the interquartile range, and points represent outlier values.

### Prediction of neurocognitive abilities

Finally, we asked whether we could use the different types of information represented by different connectome scales to tease apart individual differences in neurocognitive performance. As a proof-of-principle analysis, we looked at whether the variation in reported heritability between general cognitive ability and learning/memory relates to the functional significance of our heritability and reliability findings. We focused on the composite scores for general cognitive ability and learning/memory based on evidence of their differential heritability (Bouchard, 2004; Plomin and Spinath, 2004; Fletcher et al., 2014; Mollon et al., 2021). An initial h^2^ estimation based on intraclass correlation (Falconer’s formula (Falconer, 1996)) yielded h^2^ of general cognitive ability = 0.39 and h^2^ of learning/memory = 0.19 (Figure 6A) for our sample, which converged with our expectations based on prior literature (Plomin and Spinath, 2004; Need and Goldstein, 2009; Fletcher et al., 2014; Mollon et al., 2021). A more sensitive heritability analysis including significant covariates for site, sex, and ethnicity, estimated h^2^ of these traits in the present cohort to be higher and more similar than the values reported in the literature: for the participants in our twin sample (*n* = 748 with complete data), we estimated h^2^ of general cognitive ability as 0.884 ± 0.01 and h^2^ of learning/memory to be 0.837 *±* 0.01 (Figure 6B). To assess whether our higher heritability scores were attributable to subject selection based on quality and amount of neuroimaging data, we replicated this analysis on the entire cohort (*n* = 9,519 with complete data) and found the same pattern of results: general cognitive ability h^2^ = 0.83 *±* 0.02, learning/memory h^2^ = 0.80 *±* 0.02 (Figure 6C).

**Figure 6:**
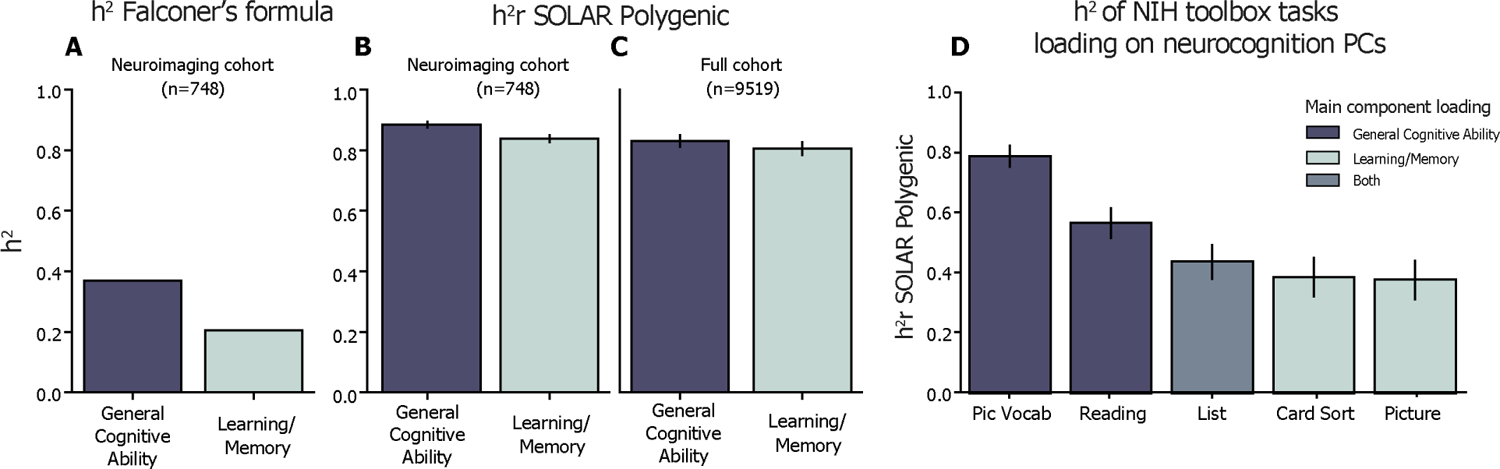
Heritability of neurocognitive measures. A. h^2^ of general cognitive ability and learning/memory were computed using Falconer’s formula, which is 2 times the difference between ICC among MZ pairs and ICC DZ pairs. h^2^ general cognitive ability = 0.39, h^2^ learning/memory = 0.19. B. h^2^r of general cognitive ability and learning/memory were computed using SOLAR polygenic across participants in our twin cohort (full sample: *n* = 778; with complete data: *n* = 748). h^2^ of general cognitive ability = 0.884 ± 0.01; h^2^ of learning/memory = 0.837 ± 0.01 (reported as SOLAR polygenic h^2^r ± standard error).

A final consideration was the possibility that the neurocognition PC loadings are a more stable phenotype than individual cognitive task scores, but that the individual tasks are more variably heritable. To investigate this, we broke down the composite scores into the tasks that primarily load onto each component in the BPPCA solution and found a scaling of heritability relative to the component score.

Tasks loading more strongly onto the general cognitive ability component (*PicVocab, Reading*) showed stronger heritability whereas tasks loading more strongly onto the learning/memory component (*CardSort, Picture*) showed weaker heritability. A task loading onto both components evenly (*List*) showed intermediate heritability (Figure 6D), which indicates that general cognitive ability and learning/memory have different genetic contributions in this sample and the composite scores are complicated by their overlapping tasks. Notably, even with overlap in the two components (i.e., *List* task loading on both), we still see distinct brain-based predictions associated with heritability, supporting our approach as a proof-of-principle analysis. Based on the neurocognitive component scores, we examined if the different types of information represented at different connectome scales captured dissociable variance in neurocognitive traits. Given that CHA improves reliability of individual differences in the fine-scale connectome, we hypothesized that CHA fine connectomes would capture more variance in both cognitive traits (general cognitive ability, learning/memory) than the CHA coarse, AA coarse, or AA fine connectomes. Moreover, we hypothesized that the most heritable connectome (AA coarse) would reflect more variance in a more heritable cognitive trait (general cognitive ability) relative to a less heritable one (learning/memory).

Significant covariates included site, ethnicity, and sex. C. To evaluate whether the neurocognitive heritability effect was due to sampling bias toward the subjects included in the neuroimaging analysis, we recalculated h^2^r (as in B) of the same components using the full ABCD study sample (full sample: *n* = 11,875; with complete data *n* = 9,519). Estimate of general cognitive ability = 0.830 ± 0.02, learning/memory = 0.800 ± 0.02. D. h^2^r analysis of the individual NIH Toolbox tasks loading onto the general cognitive ability and learning/memory components. Tasks loading more strongly onto the general cognitive ability component (*PicVocab, Reading*) showed stronger heritability whereas tasks loading more strongly onto the learning/memory component (*CardSort, Picture*) showed weaker heritability and a task loading onto both components evenly (*List*) showed intermediate heritability. Abbreviations: PicVocab = Toolbox Picture Vocabulary Task (language skills and verbal intellect); Reading = Toolbox Oral Reading Recognition Task (read and pronounce single words), List = Toolbox List Sorting Working Memory Test; CardSort = Toolbox Dimensional Change Card Sort Task (cognitive flexibility); Picture = Toolbox Picture Sequence Memory Test.

We used a principal components analysis (PCA) ridge regression to predict scores on these two cognitive domains from connectivity profiles at each scale. Models were trained using the individual differences matrices (IDMs) across 526 unrelated subjects, computed for each scale, alignment, and parcel. Crucially, predictions based on IDMs ensure that all models are receiving data with the same number of features, regardless of connectome type, since IDMs are 526 subjects by 526 subjects where each cell is the pairwise covariance of the connectomes of two unrelated subjects. We used a nested cross-validation procedure to tune hyperparameters (number of PCs and α) then applied the model to predict loadings from unseen subjects (see *Methods: Neurocognition PC prediction analysis* for more details). Regression model scores are presented as *R*^2^ between predicted scores and true values, p-values for each parcel are assessed by permuting neurocognition scores 1,000 times, and scores were compared across connectome types using permutation tests with 10,000 iterations.

General cognitive ability was similarly predicted by CHA fine, AA coarse, and AA fine connectomes (average *R*^2^ *± s*.*d*. across parcels: CHA fine = 0.0133 *±* 0.021; AA fine = 0.0134 *±* 0.020; AA coarse = 0.0124 *±* 0.018; Bonferroni-corrected p-values: CHA fine vs. AA fine *p_bonf_* = 1.0; CHA fine vs. AA coarse *p_bonf_* = 1.0; AA fine vs. AA coarse *p_bonf_*= 0.39) (Figure 7 left), indicating that the information content carried in each of these connectomes predicts general cognitive ability. Compared with the other three connectomes, CHA coarse connectomes predicted significantly less variance in general cognitive ability (*R*^2^ = 0.006 *±* 0.02; CHA coarse vs. CHA fine *p_bonf_ <* 0.0001; CHA coarse vs.

**Figure 7:**
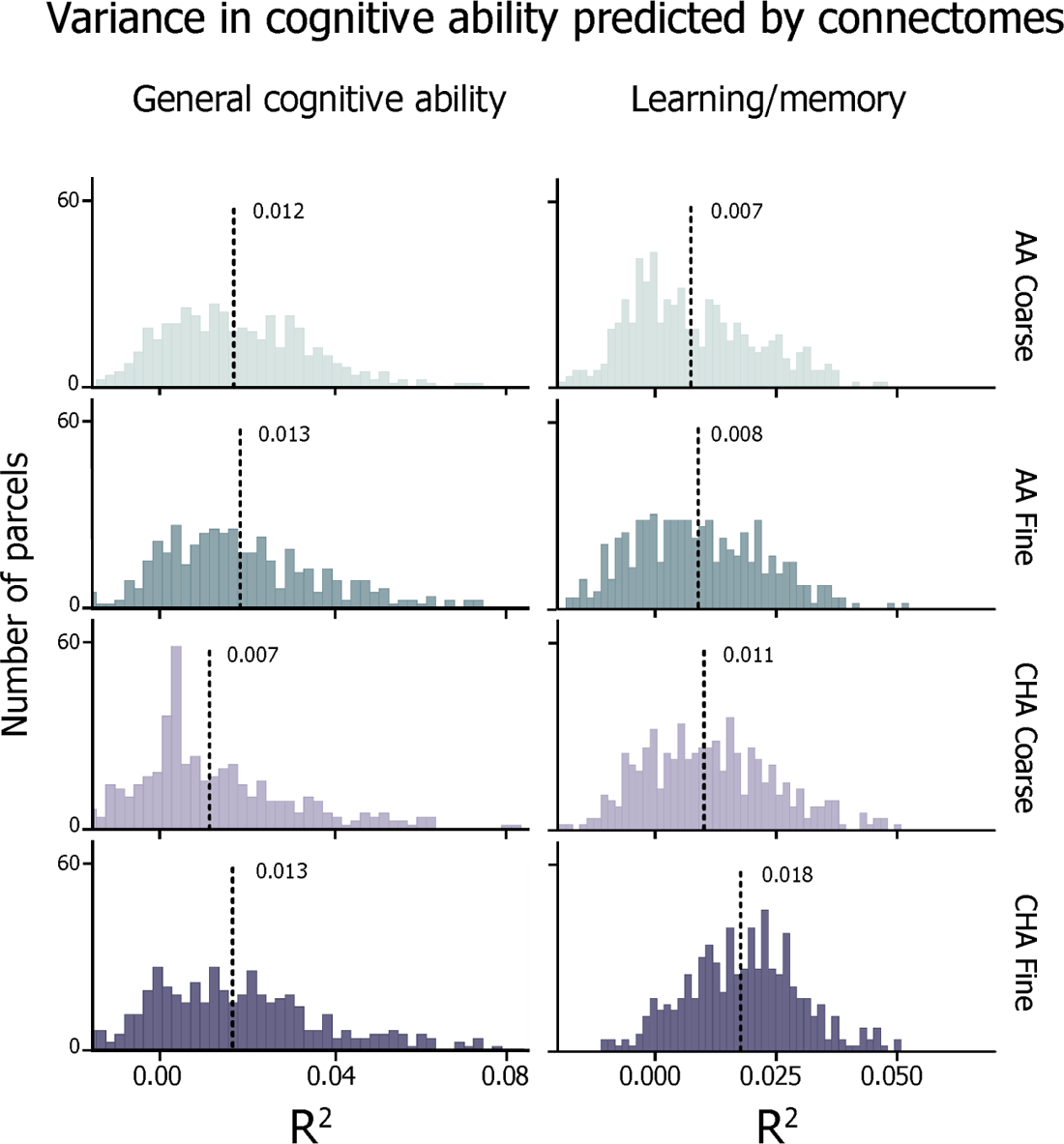
Neurocognition prediction results. Distribution of *R*^2^ values across all parcels for prediction of general cognitive ability (left) and learning/memory (right). Histogram heights correspond to density of scores and lines represent the mean of the score distribution. For the purpose of distribution matching, *R*^2^ values are not thresholded here.

AA fine *p_bonf_ <* 0.0001; CHA coarse vs. AA coarse *p_bonf_ <* 0.0001), suggesting that functionally aligning fine-scale information then smoothing over the functional information space actually diminishes the specificity of the information. After performing permutation tests at each parcel and thresholding at *p_FDR_ <* 0.05, 52.7%, 53.3%, 35%, and 54% of parcels predicted general cognitive ability significantly greater than chance at the AA coarse, AA fine, CHA coarse, and CHA fine, respectively.

Learning/memory is regarded as a less heritable psychological trait (h^2^ = 0.36 – 0.56) (Mollon et al., 2021), so we hypothesized that the information content of the coarse connectomes (the more heritable information) would contribute less to the prediction of this trait relative to the fine connectomes (the more reliably idiosyncratic information). Prediction of learning/memory scores were significantly higher for the CHA fine connectome than the other connectome types (average *R*^2^ *± s*.*d*. across parcels: CHA fine = 0.018 *±* 0.012; CHA coarse = 0.011 *±* 0.014; AA fine = 0.008 *±* 0.014; AA coarse = 0.007 *±* 0.014; Bonferroni-corrected p-values: CHA fine vs. CHA coarse *p_bonf_ <* 0.0001; CHA fine vs. AA fine *p_bonf_ <* 0.0001; CHA fine vs. AA coarse *p_bonf_ <* 0.0001; CHA coarse vs. AA fine *p_bonf_*= 0.007; CHA coarse vs. AA coarse *p_bonf_ <* 0.0001; AA fine vs. AA coarse *p_bonf_* = 0.529) (Figure 7 right). The gap in the percentage of parcels capturing a significant amount of variance was also more pronounced for this trait; 37.2%, 40%, 53.1%, and 80.5% for AA coarse, AA fine, CHA coarse, and CHA fine, respectively.

Prior work has indicated that in-scanner head movement is correlated with neurocognition scores. Despite strict participant exclusion based on data quality control (see Figure 1), our cohort still showed a mild relationship between mean framewise displacement (FD) and neurocognitive scores (*n* = 526; Pearson’s *r* = *−*0.073, *p* = 0.09 for general cognitive ability; *r* = *−*0.135, *p <* 0.05 for learning/memory). To control for possible effects of movement on our prediction, we repeated the neurocognition PC prediction analysis controlling for FD. Model performance after controlling for FD was almost identical to our original results; the correlation of model performance between FD-controlled prediction and original prediction was *r* = 0.999 for general cognitive ability and *r* = 0.995 for learning/memory (Figure 8).

**Figure 8:**
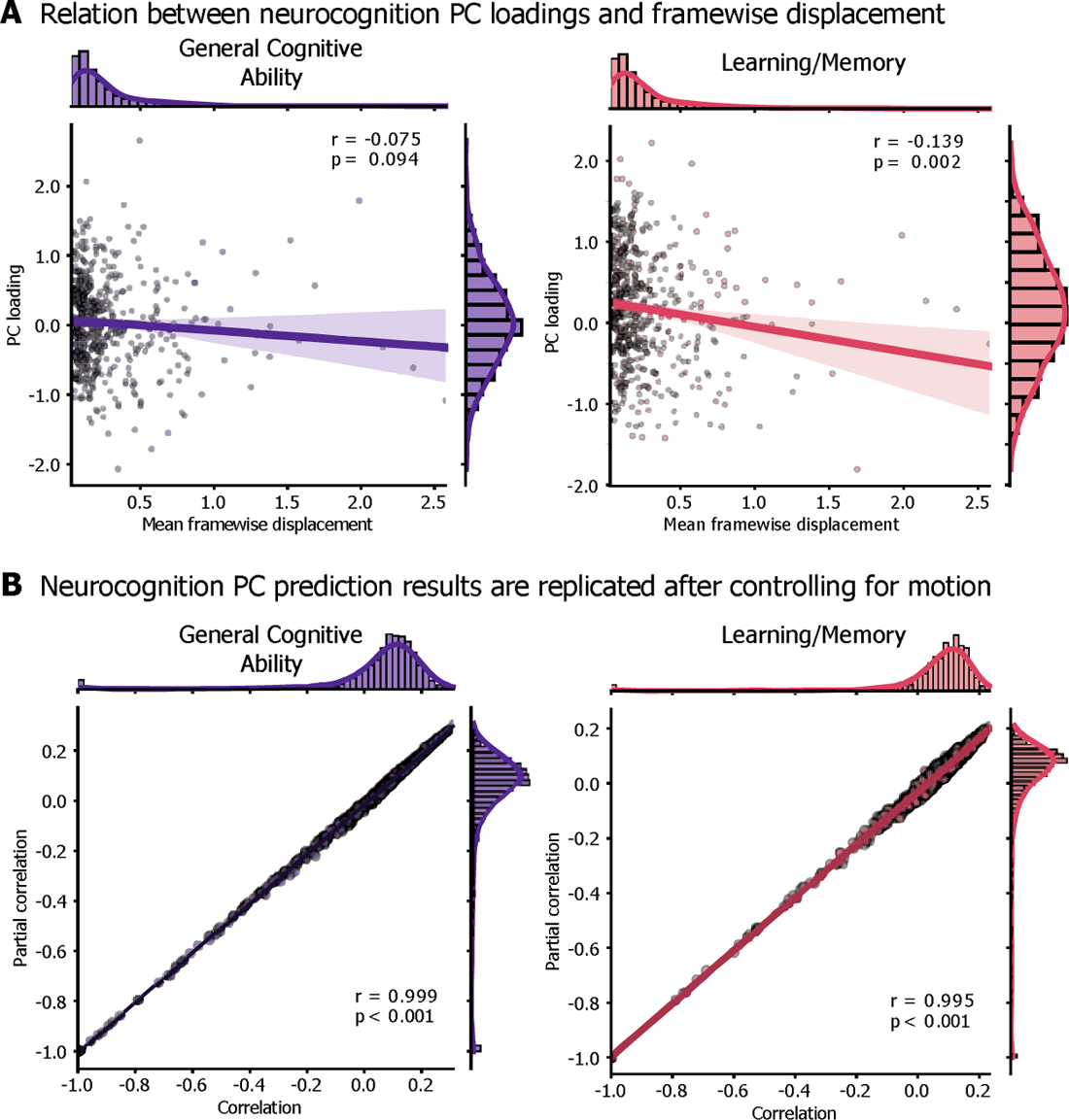
Replication of prediction results after controlling for head movement. A. Participants’ loadings onto the neurocognition PCs are mildly anti-correlated with mean framewise displacement. Each participant included in the neurocognition PC analysis (502 with complete data) is represented as a point in the scatterplots. B. To account for the relationship between head movement and cognitive scores, we replicated the prediction analysis controlling for mean framewise displacement. After getting predicted scores for each participant with leave-one-out cross validation, we then either computed Pearson’s correlation between model predictions and true scores (“correlation” on the x-axis) or a partial correlation, including framewise displacement as a covariate (“Partial correlation” on the y-axis).

Prediction results are replicated after controlling for movement. Note that in main analysis (Figure 7), the prediction results (without the additional control for movement) are presented as adjusted *R*^2^, but for equal comparison with the partial correlation, we present prediction results as a Pearson’s correlation, prior to the adjusted *R*^2^ calculation.

## Discussion

In this study, we investigated whether heritable and experiential factors have dissociable effects on the development of coarse- and fine-grained functional connectomes. We hypothesized that the coarse structure of connectivity between cortical regions would be under stronger genetic control, whereas the fine structure of local variations in connectivity would show more pronounced and reliable individual differences. We improved access to fine-scale connectivity by applying connectivity hyperalignment (CHA) to resting-state fMRI data in a large sample of children. Our study presents the first example of defining and applying a fine-scale functional common space to a developmental cohort. This represents an important advancement as it disentangles shared connectivity patterns and meaningful individual differences from misaligned cortical anatomy or measurement noise, which commonly plague developmental neuroimaging (Power et al., 2012; Satterthwaite et al., 2012; Birn et al., 2013; Fair et al., 2020; Kennedy et al., 2022).

Using RSFC from a representative sample of 200 unrelated children, we built a fine-scale model connectome and hyperaligned a separate set of 915 children’s data to that model. Pre-hyperalignment, anatomically aligned connectivity patterns exist in distinct, individual representational spaces and are, therefore, both less reliable and less predictive of cognition. Hyperalignment resamples an individual’s vertex-wise connectivity profiles into the model connectivity profiles, which affords meaningful comparisons of vertex-wise profiles across individuals. Prior literature has relied upon anatomical alignment, or matching children’s data to a common anatomical template, then applying a parcellation to extract one coarse pattern per parcel (Marek et al., 2019, 2022; Sripada et al., 2020; Chen et al., 2022). Averaging connectivity profiles across neighboring vertices within a parcel can factor out misaligned vertices to reveal coarse connectivity structure, but obscures fine-scale information. We considered multiple connectome types — coarse- and fine-grained, pre- and post-CHA — to understand the effects of genetics and experience at different scales of connectivity.

Using CHA, we revealed shared patterns of fine-scale FC with reliable individual variation. The coarse-scale connectome (AA coarse) shows only moderately reliable idiosyncrasies (mean *r* = 0.34), versus the CHA fine connectome’s mean reliability of *r* = 0.86 (Figure 4). Across the cortex, individual differences in fine-scale FC were more reliable than the coarse-scale, suggesting that traditional alignment diminishes spatially resolved functional topographies. By estimating the heritability of RSFC at each scale and alignment in twins, we showed that idiosyncrasies in fine-scale connectomes were dissociable from those in the coarse-scale connectome. Multiscale heritability analysis of AA coarse connectomes attributed about 20% (5.47–33.06%) of variance to genetic relatedness, converging with expectations based on adult RSFC (Glahn et al., 2010; Miranda-Dominguez et al., 2018; Anderson et al., 2021) and how neuroimaging phenotypes show increasing heritability with age (Blokland et al., 2012; Elliott et al., 2018). At the fine scale, h^2^ estimates dropped to around 10% regardless of alignment (CHA: 3.1–19.2%, AA: 2.0–16.67%), suggesting that idiosyncrasies in fine-scale RSFC were likely more associated with unique experience and gene–environment interactions than genetic factors (Figure 5). Individual differences in fine-scale connectivity after hyperalignment were *more* reliable than differences in coarse-scale connectivity, but they were *less* heritable, indicating a dissociation between the spatial scales of heritable and reliable RSFC. Our final analysis examined the cognitive relevance of this dissociation. The most reliably idiosyncratic (CHA fine) and most heritable (AA coarse) connectomes similarly predicted general cognitive ability, a canonically heritable cognitive trait. In contrast, the most reliably idiosyncratic connectome (CHA fine) best predicted a less heritable cognitive trait (learning/memory) (Figure 7).

Taken together, we demonstrate a link between reliable, heritable, and predictive scales of FC and highlight the importance of nuanced, multi-scale connectivity analyses for comprehensively understanding individual differences during development. Our findings dovetail with the interactive specialization framework of functional brain development, which states that the development of the functional connectome is not solely determined by a blueprint, but the interaction of a heritable template of connections and individualized experience (Johnson, 2011). We present neural evidence of this framework: the coarse connectome serves as a heritable blueprint which represents more heritable cognitive phenotypes. The blueprint encoded in coarse-scale connectomes becomes part of the fine-scale connectome during development, as prediction models based on AA coarse never outperform those based on CHA fine, but the fine-scale connectome affords cognitive prediction beyond the coarse.

Recent work has recognized the need for massive numbers of subjects to reliably associate brain phenotypes with behavior, specifically in developmental RSFC (Marek et al., 2022). This could be attributed to the complexity of neuroimaging phenotypes, but by leveraging machine learning advances like multivariate models, functional alignment, and individualized atlases, we can improve prediction (Dubois and Adolphs, 2016; Kong et al., 2019, 2021, 2023; Rosenberg et al., 2020; Feilong et al., 2021, 2022; Chen et al., 2022; DeYoung et al., 2022). We demonstrate hyperalignment’s power as a tool for understanding developing brain–behavior associations. Here, hyperalignment was trained on RSFC from 200 children and tested on a separate cohort, using <20 minutes of rs-fMRI per subject. The prediction results assure that the increased reliability of individual differences was driven by generalizable, shared signals in the hyperalignment model rather than overfitting to subject-specific signals or simply higher-dimensional data, since CHA improved reliability over AA at the fine-scale as well. By performing subsequent analyses on (dis)similarity matrices, we ascertain that our findings are driven by the information content represented in different connectomes, rather than the dimensionality difference between coarse- and fine-scale connectomes.

This study should be interpreted in light of potential limitations. Though statistically significant, our predictions accounted for low variance in cognitive measures relative to adult studies (Finn et al., 2015; Dubois et al., 2018b; Feilong et al., 2021). While task-based FC is more stable and behaviorally predictive than RSFC among adults (Rosenberg et al., 2016; Greene et al., 2018; Feilong et al., 2021), this requires sufficient amounts of high-quality data challenging to collect in children (Satterthwaite et al., 2012; Kennedy et al., 2022). Prior work shows that RSFC is more heritable than task-based FC (Elliott et al., 2018), suggesting that tasks add cognitive load and introduce idiosyncratic information processing less attributable to genetic factors.

Our focus on neurocognition composite scores should be viewed as a proof-of-principle analysis relating the heritability and reliability of individual differences in neuroimaging phenotypes to differentially heritable behavioral phenotypes due to limitations in h^2^ estimations of psychological traits. An initial model of neurocognitive trait heritability (ICC-based Falconer’s formula) yielded estimates of general cognitive ability and learning/memory convergent with prior literature (Plomin and Spinath, 2004; Need and Goldstein, 2009; Fletcher et al., 2014; Mollon et al., 2021). A more sensitive model (SOLAR Polygenic) including demographic covariates yielded h^2^ estimates higher and more similar than previously reported. Possible explanations for this discrepancy include our sample’s demographic differences with the literature: our sample is aged 9–10 years, whereas prior studies report larger age and h^2^ ranges (h^2^ of general cognitive ability = 0.54−0.85 at 10–12 years (Bouchard, 2004), 0.60−0.73 at 8–21 years (Mollon et al., 2021); h^2^ of learning/memory = 0.18 − 0.55 at 8–21 years (Mollon et al., 2021) and 0.39 at 18–67 years (Fletcher et al., 2014)). Our sample is more racially diverse than prior studies, which generally control for genetic background by using homogeneous samples, predominantly individuals of European descent (Glahn et al., 2010; Haworth et al., 2010; Popejoy and Fullerton, 2016; Savage et al., 2018; Grasby et al., 2020). Our predictions align with the h^2^ values reported by Mollon et al. (2021), which used a younger and more diverse cohort than prior studies and more like our sample. Future work could investigate whether the heritability scores become clearer with age and how that relates to brain-based prediction, using a future release of ABCD data.

This paper presents several novel findings. First, hyperalignment can be used as a tool to improve the reliability and behavioral relevance of functional connectivity in children. After hyperalignment, reliability of individual differences in RSFC is improved at the fine-scale relative to the coarse scale, but the heritability of these signals is weakened. Finally, we present a potential application of this dissociation in the fine and coarse scale connectomes to show how the heritability of RSFC relates to ability to predict cognition based on the heritability of neurocognitive traits. With the substantial twin cohort, behavioral testing, and longitudinal neuroimaging data included in the ABCD study, future work could investigate the connection between heritability and predictivity of functional connectivity over time, as the heritability of both neurocognitive measures and neuroimaging-based phenotypes increase with age (Thompson et al., 2001; Haworth et al., 2010; Blokland et al., 2012; Jansen et al., 2015). Moreover, this approach could be used to investigate how functional topographies reflect cognitive development over time, by hyperaligning a subject to themselves at multiple timepoints. In conclusion, the current study shows that by breaking apart the pieces of the functional connectome into their coarse and fine structures, we can better understand how these scales interact to scaffold and instantiate reliable idiosyncrasies in brain and cognition during development.

## Author contributions

Conceptualization, ELB, KMR, MDR, MF, JVLH, RW, and BJC; Methodology, ELB, MF, JVLH, and KMA; Software, ELB, KMA, and MF; Validation, ELB, KMA, MF, and JVLH; Investigation, ELB; Writing - Original Draft, ELB, MF, and BJC; Writing - Review & Editing, ELB, KMR, MDR, RW, KMA, MF, JVH, and BJC; Visualization, ELB and KMR; Supervision, MF, JVH, and BJC; Funding acquisition, BJC.

## Notes

### Competing Interest Statement

Kevin Anderson is an employee at Neumora Therapeutics.

### Summary of Updates

Updated with respect to reviewer comments; methods and discussion clarified.

